# Role of toll-like receptor 2 during infection of *Leptospira* spp.: A systematic review

**DOI:** 10.1101/2023.08.23.554396

**Authors:** Chamila Niroshani Kappagoda, Indika Senavirathna, Suneth Agampodi

**Affiliations:** Department of Community Medicine, Faculty of Medicine and Allied Sciences, Rajarata University of Sri Lanka, Saliyapura, Anuradhapura, Sri Lanka; Department of Biochemistry, Faculty of Medicine and Allied Sciences, Rajarata University of Sri Lanka, Saliyapura, Anuradhapura, Sri Lanka; Department of Internal Medicine, Section of Infectious Diseases, School of Medicine, Yale University, New Haven, Connecticut, United States of America; International Vaccine Institute, Seoul, Republic of Korea

## Abstract

**Background:** Present systematic review was conducted to determine the role of the Toll-like receptor 2 during *Leptospira* infection in *in-vitro*, *in-vivo*, and *ex-vivo* experimental models and human studies.

**Methods:** Original articles published in English up to March 2022 that examined the response of Toll-like receptor 2 during leptospirosis were selected. PubMed, Web of Science, Scopus, Trip, and Google Scholar were used to search the literature. The National Institute of Health Quality Assessment tool, Systematic Review Centre for Laboratory Animal Experimentation risk of bias tool, and Office of Health Assessment and Translation extended tool were used to assess the risk of bias and the quality of the studies.

**Results:** Out of 2406 studies, only 32 were selected for the systematic review. These comprised 3 human studies, 14 *in-vitro* studies, 5 *in-vivo* studies, and 3 *ex-vivo* studies. 7 studies employed combined models that encompassed human, in-vivo, in-vitro, and ex-vivo. In our analysis, we assessed the response of Toll-like receptor 2 (TLR2) through various indicators, including TLR2 receptor/mRNA expression and indirect TLR2 involvement via the secretion/mRNA expression of cytokines, chemokines, and immune effectors. Notably, we identified increased TLR2 expression and the secretion/mRNA expression of several cytokines (IL6, IL8, IL-1β, TNFα, IFNγ, IL10, CCL2/MCP-1, CCL10, COX2, CXCL1/KC, CXCL2/MIP2) and immune effectors (hBD2, iNOS, Fibronectin, Oxygen, and Nitrogen reactive species) as key aspects of host TLR2 responses during leptospirosis. Besides the role of TLR2 in response to leptospirosis, the involvement of TLR4 and TLR5 was identified in *in-vitro* and *in-vivo* studies. IL6, IL10, IL-1β, TNFα, MIP, CCL2, CCL10, COX2, MCP1, IFNγ, iNOS, NO, anti-*Leptospira* IgG were triggered through TLR4. Furthermore, TNFα secretion was stimulated through TLR5. In addition to the role of TLR2, our review revealed the involvement of TLR4 and TLR5 in in-vitro and in-vivo studies. Specifically, the activation of TLR4 triggered responses including IL6, IL10, IL-1β, TNFα, MIP, CCL2, CCL10, COX2, MCP1, IFNγ, iNOS, NO, and anti-Leptospira IgG.

**Discussion:** Recognition of pathogen-associated molecular patterns through TLR2 triggers the secretion of cytokines/chemokines and immune mediators, facilitating the eradication of *Leptospira* infection. However, excessive amounts of these compounds can harm host tissues; therefore, regulating immune mediators through TLR2 using agonists or antagonists at an optimal level is important for mitigating tissue damage and promoting effective immune responses. In addition to TLR2, TLR4 and TLR5 were found to play defensive roles in *in-vitro* and *in-vivo* studies against *Leptospira* infection.

**Other:** *Funding:* No funding received for this study.

*Registration:* PROSPERO 2022 CRD42022307480

**Author summary:** Leptospirosis is a globally widespread, infectious zoonosis caused by a spiral shape bacterium belonging to the genus *Leptospira*. Pathogenic *Leptospira* spp. play a significant role in infecting humans resulting in a wide range of clinical symptoms ranging from febrile illness to multi-organ failures. Different host immune responses are the key contributors to the disease development, pathogenesis factors of the infectious organism, and epidemiological factors. Host immune responses initiate by interacting with the pathogen’s molecular patterns and the host immune cell receptors. In global literature, Toll-like receptors are the mainly studied host pattern recognition receptors, with Toll-like receptor 2 plays a crucial role in mediating the human immune responses. Although there are narrative reviews regarding the role of Toll-like receptor 2, it is worth systematically reviewing it with methodological rigor. The secretion of the cytokines/chemokine and immune mediators will facilitate the elimination of bacterial infection. However, excessive amounts of these compounds can harm host tissues; therefore, regulating immune mediators through Toll-like receptor 2 using agonists or antagonists at an optimal level is essential. Despite the disease burden, the lack of advanced treatments and efficient diagnostic methods hinders disease management. Exploring host immune responses against the disease through Toll-like receptor 2 could provide valuable insights for the development of therapeutic strategies.

## Introduction

Human innate immune responses to pathogens are triggered through the recognition of Pathogen Associated Molecular Patterns (PAMPs) by various human Pattern Recognition Receptors (PRRs). PAMPs are expressed in the outer cell membrane of the pathogens, which is essential for their survival and virulence. Lipopolysaccharides (LPS) and proteins are the major PAMPs of pathogenic *Leptospira* spp.[1–4], which are obligate aerobic spirochetes that cause leptospirosis. Although a variety of PRRs are possessed by innate immune cells during infection with *Leptospira* spp., Toll-like receptors (TLRs) are among the most studied PRRs in infectious disease research[5]. The ‘Toll’ gene of *Drosophila melanogaster* (Common fruit fly) was first isolated and characterized by Carl Hashimoto and colleagues in 1988 [6]. The first TLR identified in mammalian tissues was TLR4 which was previously named as hToll. Gene expression of TLR4 has been reported in monocytes, macrophages, dendritic cells, γδ Tcells, and small intestinal cell lines of humans and mice [7]. Since the identification of TLR4 in mammals, 13 TLRs (TLR1-TLR13) have been discovered and described in the literature [8]. Numerous *in-vitro* and mice model studies have been conducted on immune responses against *Leptospira* spp., and TLR1, TLR2, TLR4, and TLR6 are considered to have direct involvement in the immune response against *Leptospira* spp. While mice serve as reservoir hosts for pathogenic *Leptospira* spp., it can be recognized through mice TLR4 but not through human TLR4[9,10].

Wertz and colleagues have shown that TLR2 of mice and human monocytic cells can trigger innate immune response via the recognition of *L.interrogans* LipL32, the major outer membrane protein in pathogenic *Leptospira* and LPS, with the presence of CD14 factor [11]. Further studies have used atomic force microscopes to assess the direct interaction of TLR2 with the cell surface receptors of the bacterium. It has provided direct evidence of TLR2 interaction with LipL32 receptors expressed on the *Leptospira* cell surface [12]. Interaction of pathogenic *Leptospira* spp. with human PRRs responds to pathogens via secretion of immune mediators, cytokines, and antimicrobial peptides and inducing the recruitment of other leukocytes[13,14] to kill the bacteria.

The response of the TLR2 was studied majorly in *in-vitro*, *in-vivo* models while scarcely studied in human studies. In this paper, we systematically reviewed the published literature to determine the role of the Toll-like receptor 2 during *Leptospira* infection in *in-vitro*, *in-vivo*, *ex-vivo* experimental models and human studies.

## Methods

The systematic review was carried out and reported in accordance with the Cochrane guidelines and Preferred Reporting Items for Systematic Reviews and Meta-Analyses Statement 2020 (PRISMA2020) [15].

### Eligibility criteria

We selected original articles published in English up to March 2022 using the PICO approach, which involves examining the Participants, Intervention, Comparison, and Outcome. We included all human studies, animal studies, and *in-vitro* studies that examined the response of TLR2 during *Leptospira* spp. infection. We excluded papers that were only abstracts, conference papers, editorials, book chapters, or author responses. Additionally, we excluded case reports, theses, reviews, and systematic reviews.

### Information sources

The final search was conducted in March 2022, using several databases, including PubMed, Web of Science, Scopus, Trip, and Google Scholar, to find relevant articles. We used specific search strategies modified for each database to identify eligible studies. Our search strategies included MeSH terms, keywords, and other important phrases. “Toll-like receptor 2”, “TLR2”, “Leptospirosis”, “Mud fever”, “Weil’s disease” and “*Leptospira* infection” were some of the key terms used in the search strategies. Additionally, we manually checked the reference lists of included articles, reviews, and systematic reviews to find additional relevant articles. The search strategies used in the systematic review are detailed in S1.

### Selection process

CK conducted the preliminary literature search. The search results were entered into Mendeley software (Mendeley Desktop version 1.19.8). Duplicate studies were screened and removed by the ‘check for duplicates’ tool in Mendeley. Title and abstract screening were carried out independently by CK and IS, according to predefined eligibility criteria. The initial screening methodology was performed in duplicate. For the final inclusion of studies, full-text screening was conducted independently and in duplicate by CK and IS. In cases of discrepancies during the study selection, SA, served to create a consensus.

### Data extraction

Data extraction was performed in a structured Excel sheet by CK and IS independently and in duplicate. The extracted data items included Article Citation, Country of origin, Aim/objectives of the study, Study participants (Human/Animal/in-vitro/ex-vivo study), Human study (age, *Leptospira* incubation period, diagnosed method, healthy control group, type of the clinical specimen/tissue investigated, experimental method used to investigate the TLR2 response, main finding of TLR2 response), Animal study (Animal model, infection duration, experimental details that deployed to analyze the TLR2 response, main finding of TLR2 response), In-vitro/Ex-vivo (Type of the cell line, infection duration, experimental details that deployed to analyze the TLR2 response, main finding of TLR2 response). In cases of disagreement between CK and IS during data extraction, reviewers discussed the disagreement, and if necessary, a third reviewer SA, was consulted for arbitration. When necessary, authors of relevant studies were contacted to clarify and obtain data.

### Study risk of bias assessment

The risk of bias of selected studies was assessed independently by CK and IS using appropriate quality assessment tools. In cases of disagreement between the two reviewers, the disagreement was discussed, and if necessary, a third reviewer, SA, was consulted, and the disagreement was resolved. Given that the systematic review included human studies, *in-vivo* studies, *in-vitro*, and *ex-vivo* studies, different quality assessment tools were used to assess the risk of bias for each type of study. Specifically, the National Institute of Health Quality Assessment tool-Quality assessment of case-control studies was used for human studies [16], the SYstematic Review Centre for Laboratory animal Experimentation (SYRCLE’s) risk of bias tool was used for animal studies [17], and the OHAT (Office of Health Assessment and Translation) extended tool [18] was used for *in-vitro* and *ex-vivo* studies. The risk of bias information was checked and recorded for each study in a table to facilitate data analysis.

### Synthesis methods

Owing to the fact that the data for this study are presented as gene expression profile, cell receptor expression, and immune responses via TLR2 recognition, the data were not suitable for combining quantitatively. Hence the meta-analysis was not performed. A narrative synthesis was conducted with the information reported in the text and tables to summarize and explain the characteristics and findings of the included studies. We explored the relationship and findings both within and between the included studies. Eventually, the role of TLR2 during the infection of *Leptospira* spp. was summarized in the present systematic review.

### Reporting bias assessment

The RoB tool SYRCLE’s presented by SYstematic Review Centre for Laboratory animal Experimentation based on the Cochrane Collaboration Risk of bias (RoB) tool [17] was used to assess the experimental animal intervention studies. The checklist (10 questions) assesses the risk of bias in the studies and addresses biases such as selection bias, performance bias, attrition bias, detection bias, and reporting bias. For *in-vivo* studies, the risk of bias was assessed and reported via the checklist of SYRCLE’s RoB tool. OHAT extended approach to assess *in-vitro* studies has 11 questions grouped into 6 types of biases (selection, confounding, performance, attrition/exclusion, detection, and selective reporting). Reporting bias was assessed and reported via appropriate questions of OHAT RoB tool.

### Certainty Assessment

GRADE (Grading of Recommendations Assessment, Development, and Evaluation)[19] domains (imprecision, inconsistency, risk of bias, indirectness, and other) were used to assess the certainty of each important outcome and an overall judgment of whether the evidence supporting a result is of high, moderate, low, or very low certainty.

## Results

### Study selection

A total of 2406 studies were retrieved from PubMed, Web of Science, Scopus, Trip, and Google Scholar. Additionally, 23 relevant articles were identified from the reference lists of existing reviews. After removing 678 duplicate articles, the remaining studies underwent screening for eligibility. Out of the initial pool, 1611 studies were excluded due to their lack of relevance to the research question, 39 were excluded as they were reviews or systematic reviews, and 13 abstracts, case reports, and book chapters were also removed. An additional 33 studies were excluded due to outcome differences. Eventually, 32 studies met the eligibility criteria and were included in the systematic review. The study selection process was illustrated in Fig 1, adhering to the guidelines outlined in the Preferred Reporting Items for Systematic Reviews and Meta-Analyses (PRISMA) flow chart. Fig 1 PRISMA flow chart shows study selection for the systematic review.

**Fig 1.**
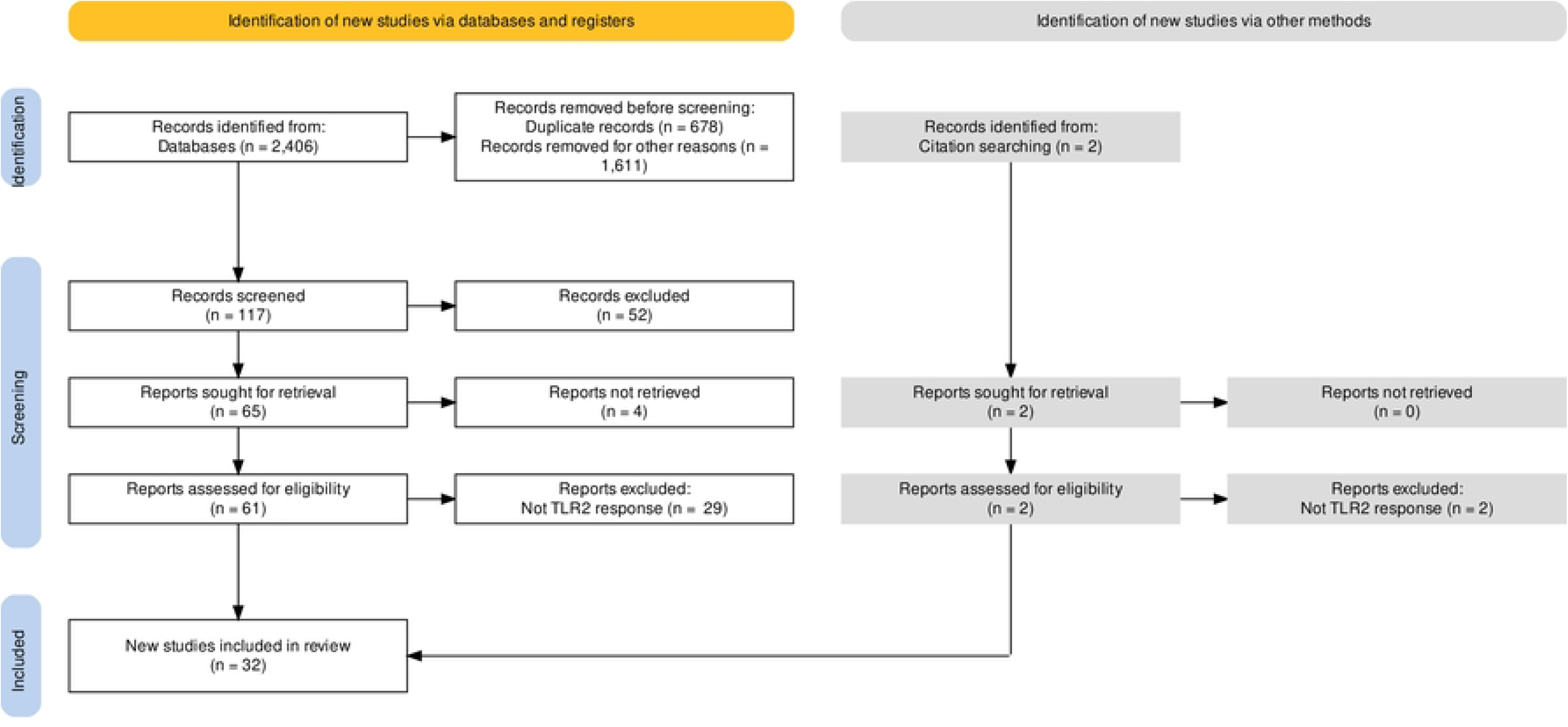
PRISMA flow chart shows study selection for the systematic review.

### General characteristics of the included studies

The 32 articles selected for this review were published between 2001 and 2021, with 7 articles each from China and Taiwan, 5 from France, 3 from the United States, 2 from Brazil, India, Thailand, and Japan, and 1 each from Argentina and the Netherlands. The studies encompassed a range of experimental models, including 3 human [20–22], 5 *in-vivo* [23–27], 14 *in-vitro* [28–41], and 3 *ex-vivo* studies [42–44]. Additionally, four studies[9,10,45,46] used a combination of *in-vivo* and *in-vitro* models, while two studies [47,48] used a combination of human, *in-vitro*, and *in-vivo* models. Finally, one study [49] used both *in-vivo* and *ex-vivo* models.

### TLR2 response in i*n-vitro* and *ex-vivo* studies

#### Direct response

Regardless of the cell line and the *Leptospira* stimulant used, TLR2 gene expression was observed as ‘increased’ compared with unstimulated cell lines [31–37,40,44]. In terms of stimulants, pathogenic *Leptospira* outer membrane proteins [40] (LipL32)[33], Loa22[31], lipopolysaccharide (LPS)[32,35–37], *L.interrogans* Copenhageni [44] stimulated the cells that increased TLR2 gene expression. Increased TLR2 gene expression was observed in mouse/rat proximal tubule cells[31,33,40], pig embryonic cells[35,36], bovine cell line[32,37], and canine whole blood[44]. In contrast, TLR2 gene expression in human kidney epithelial cells infected with *L.interrogans* Autumnalis did not differ from unstimulated cells[41]. Further, *in-vitro* studies observed TLR2 gene expression within 48hrs of infection, showing the TLR2 response in the early stage of the disease.

#### Indirect response

Blocked TLR2 / TLR2 deficient cell lines that stimulated using pathogenic *Leptospira* organism/cell components showed inhibition or decreased level of cytokine/immune mediators compared to intact TLR2 cell lines. Even with the heterogeneous stimulants and the cell lines, IL6[9,10,28,29,37,41,45,47], IL8[9,38,41–43], IL 1β[29,41,43,47,48], TNFα[9,28,29,37,38,41,43,45,47,48], IFNγ[29], IL10[46,48], iNOS[31,33], CCL2/MCP-1[29,31,33,38,39], hBD2[41,43], CCL10[29,38–40], COX2[29], CXCL1/KC[39], NFκB[48], Fibronectin[30] and CXCL2/MIP2[40] showed the inhibition or decreased level in TLR2 blocked/deficient cell lines. The data convey the involvement of TLR2 in triggering the immune response during the *Leptospira* infection. Table 1 and Table 2 show the characteristics of *in-vitro* and *ex-vivo* studies, respectively.

**Table 1.**
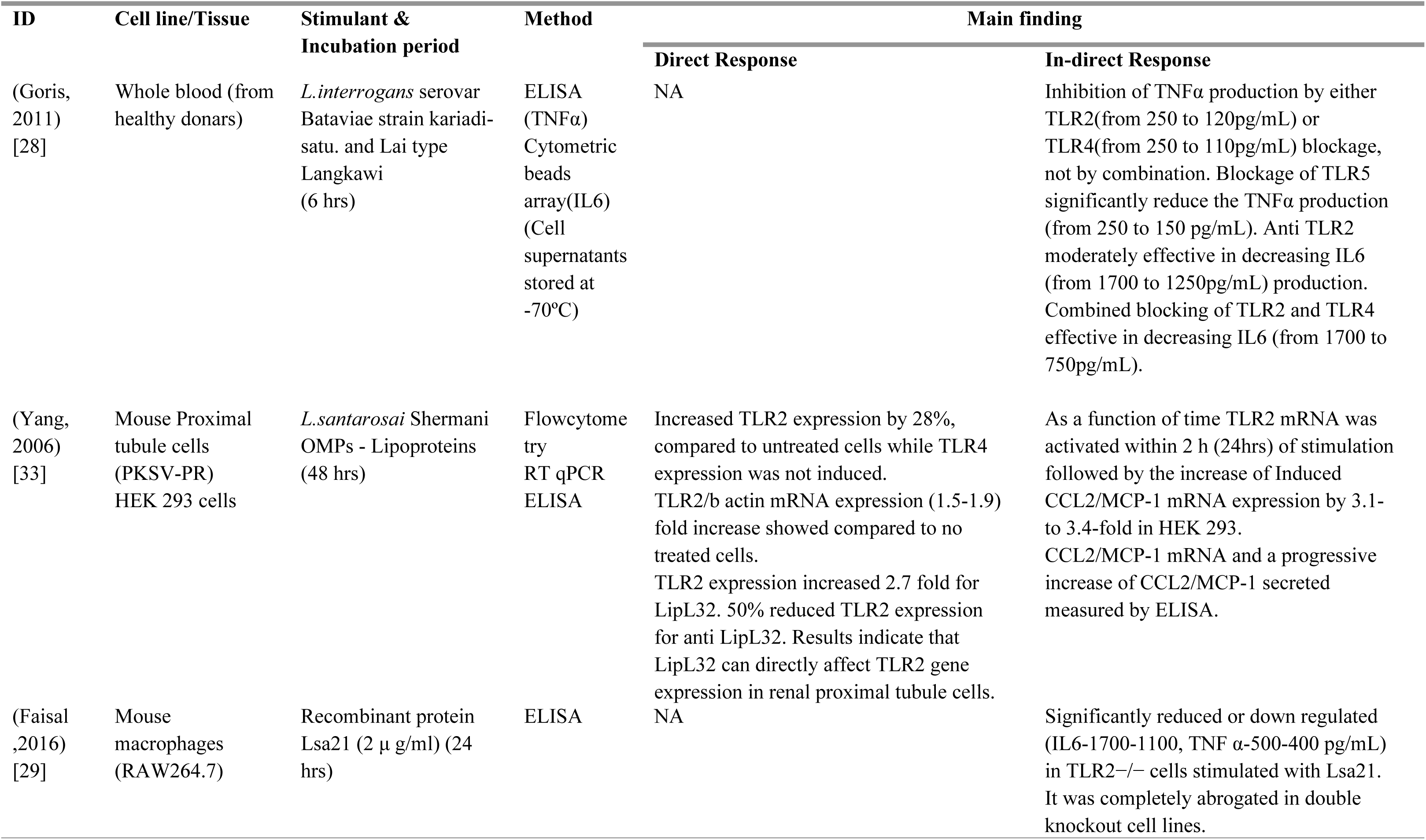
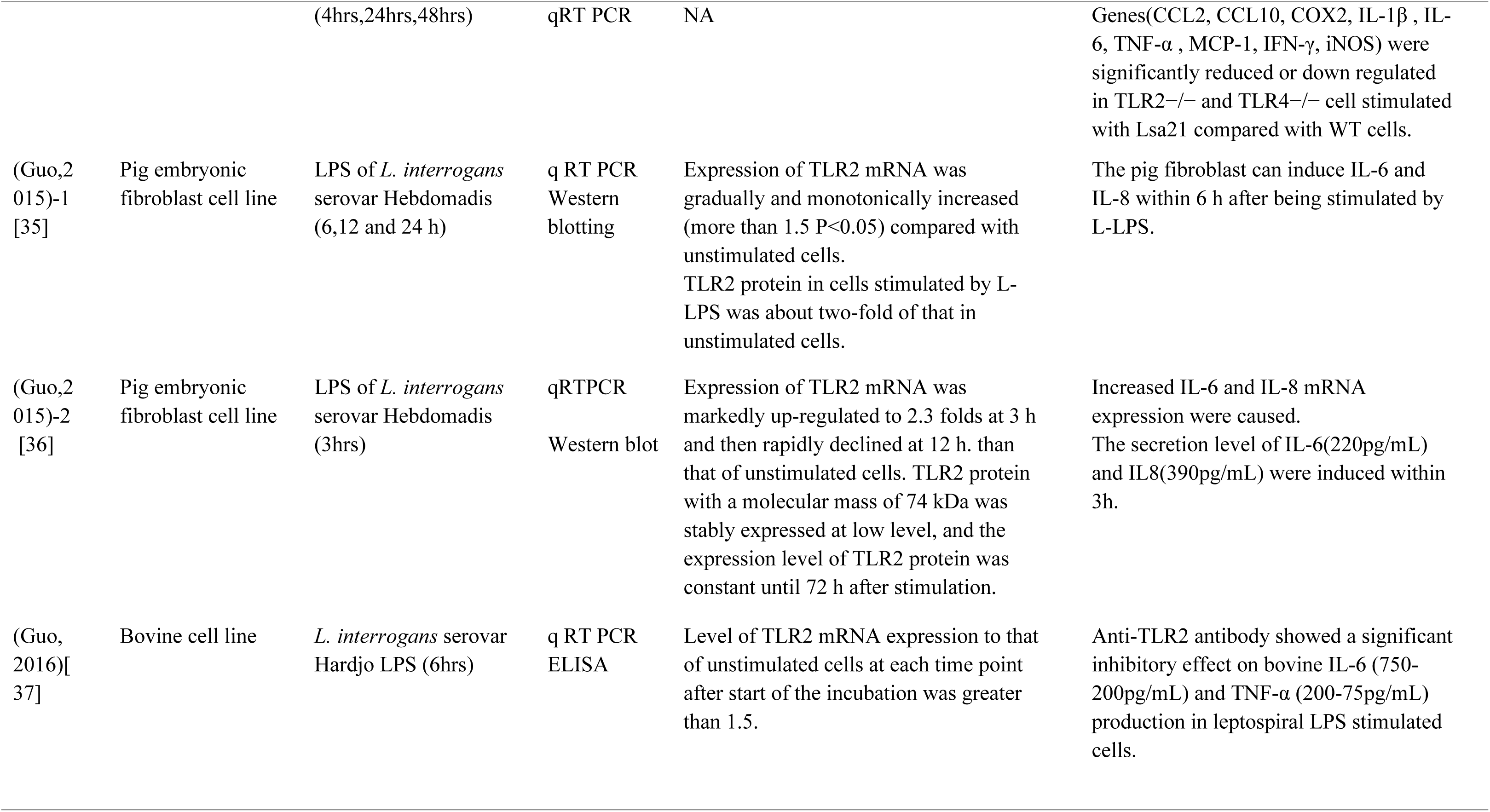
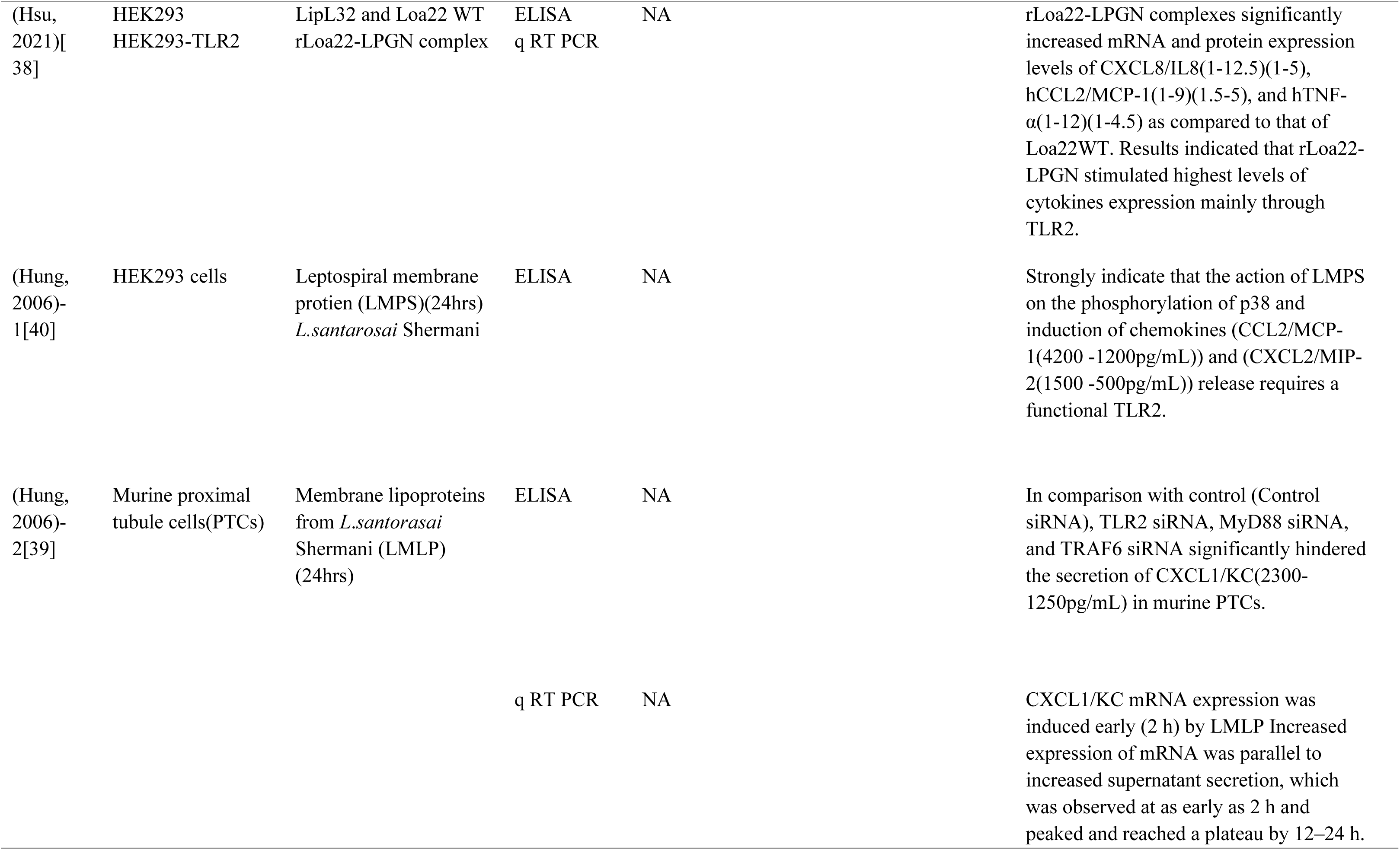
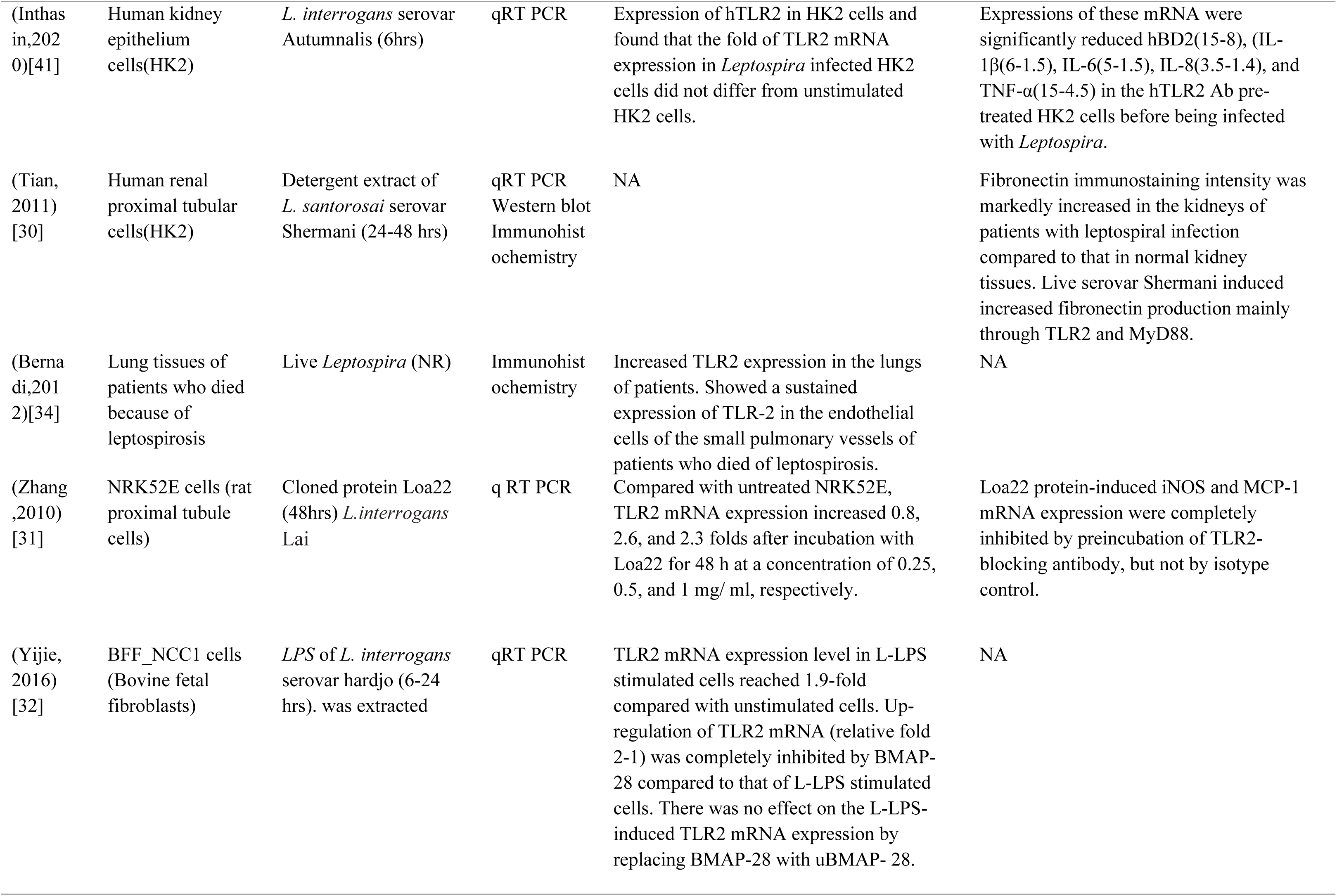
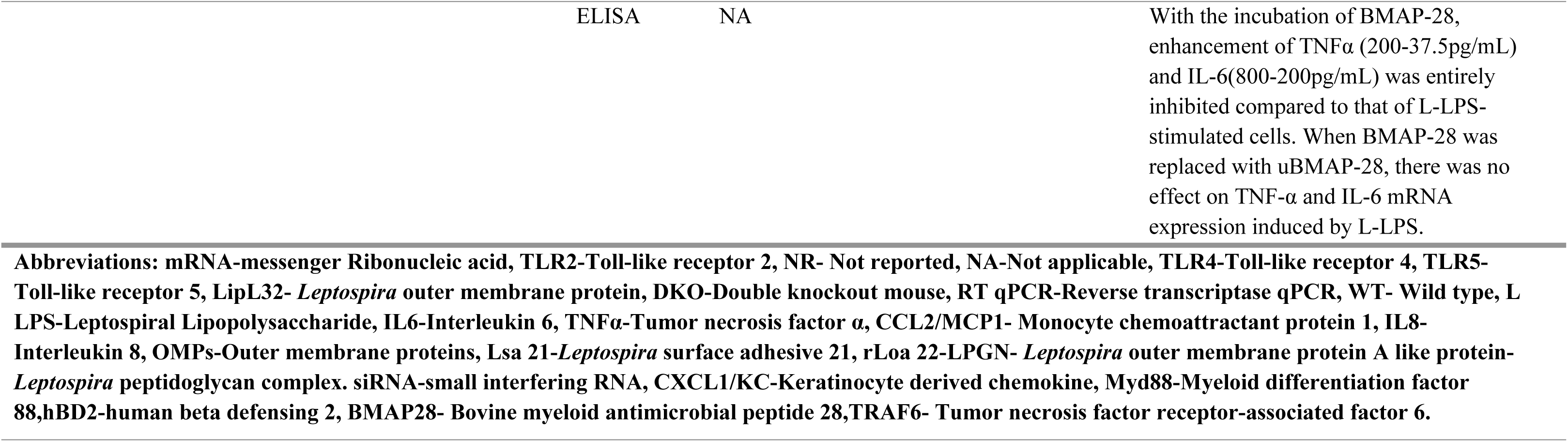
*In-vitro* study characteristics.

**Table 2.** *Ex-vivo* study characteristics.

| ID | Cell line<br>/Tissue(Ex-vivo) | Stimulant &<br>incubation period | Analysis &<br>sample storage | Main finding |  |
| --- | --- | --- | --- | --- | --- |
|  |  |  |  | Direct Response | In-direct Response |
| (Rajeev, 2020) [44] | Canine whole blood | *L. interrogans* Copenhageni<br>*L. biflexa* Patoc (18 hrs) | RT2 profiler PCR array (Cell pellets in liquid Nitrogen) | Upregulated TLR2 gene (fold change TLR2-20.30) was observed in all of the individual canine blood samples upon stimulation with *L.interrogans* and *L. biflexa* compared to controls. | NA |
| (Inthasin, 2018) [43] | Human oral tissues | *L. interrogans* serovar Autumnalis (4hrs) | RT qPCR (cDNA at -20°C) | NA | Expression (fold change) of pro-inflammatory cytokine IL-1β(8 to 3), IL-8(32 to 8), TNF-α(9 to 4), and hBD2(12 to 2) were significantly reduced in the presence of anti -hTLR2 Ab. |
| (Charo, 2019) [42] | Healthy human blood | Fiocruz L1–130 of *L. interrogans* (3-18hrs) | ELISA (NR) | NA | Increased expression of IL-8(1200 to 750 pg/mL) induced by Patoc or LIC was suppressed in neutrophils that were previously treated with an anti-TLR2, but not with an anti-TLR4. |
| Abbreviations: PCR-polymerase chain reaction, TLR2- Toll-like Receptor 2, TLR4-Toll-like Receptor 4, RT qPCR-Reverse transcriptase quantitative PCR, cDNA-complementary DNA, IL 1β- Interleukin 1β, IL8- Interleukin 8, TNFα- Tumor necrosis factor $\alpha$, hBD2-human beta defensin 2, ELISA-Enzyme linked immunosorbent assay, LIC-*L.interrogans* Copenhageni, NR-Not reported, NA-Not applicable. | | | | | |

### TLR2 response in *In-vivo* studies

#### Direct response

Despite the fact that TLR4 is the most defensive PRR in mice/hamster models[10,23] against leptospirosis, TLR2 contributed to triggering the immune response. Differential expression of TLR2 gene observed in mice renal cells upon the infection of *L.interrogans* Copenhageni[25]. Inducible effect of ‘Agonist’ components like *E.coli* LPS[26], Iris polysaccharide[27], Pam3CSK4[46] on TLR2 followed *Leptospira* infection increased the TLR2 expression in hamsters[26,27] and mice[46] models compared with infected controls. TLR2 involvement in the pathological process of *Leptospira* infection was detected in Mice[23,25] and Syrian golden hamsters[26,27]. Double knockout mice(DKO)(tlr2-/-,tlr4-/-) were very susceptible to infection, while all the wild-type mice(WT) and tlr2 −/− mice survived [23].

#### Indirect response

Mice models deficient for TLR2 showed reduced levels of TNFα, IL6[9] and reduced mRNA of IFNγ, iNOS[23] compared with wild-type mice. ‘Agonist’ activation of TLR2 increased the expression of IL1β, TNFα in mice kidney, liver, and lungs[27]. Moreover, TLR2 ‘Agonist’ Pam3CSK4 improved the ratio of IL10/ TNFα which known to be protective against infection [46]. TLR2 knockdown Zebra fish larvae demonstrated reduced kidney injury compared with wild-type zebrafish upon LipL32 infection [24]. TNFα, IL6, IL1β, IFNγ, iNOS, and IL10 expressed and/or secreted both in *in-vitro* and *in-vivo* studies with the involvement of TLR2. Diverse findings on TLR2-dependent IL6 and TNFα secretion, observed in our original study[23] and narrative review[50] that showed increased IL6 and TNFα mRNA level in DKO mice compared with WT mice. This observation implies, that other PRR apart from TLRs responsible for IL6 and TNFα production. **Error! Reference source not found.**

**Table 3.**
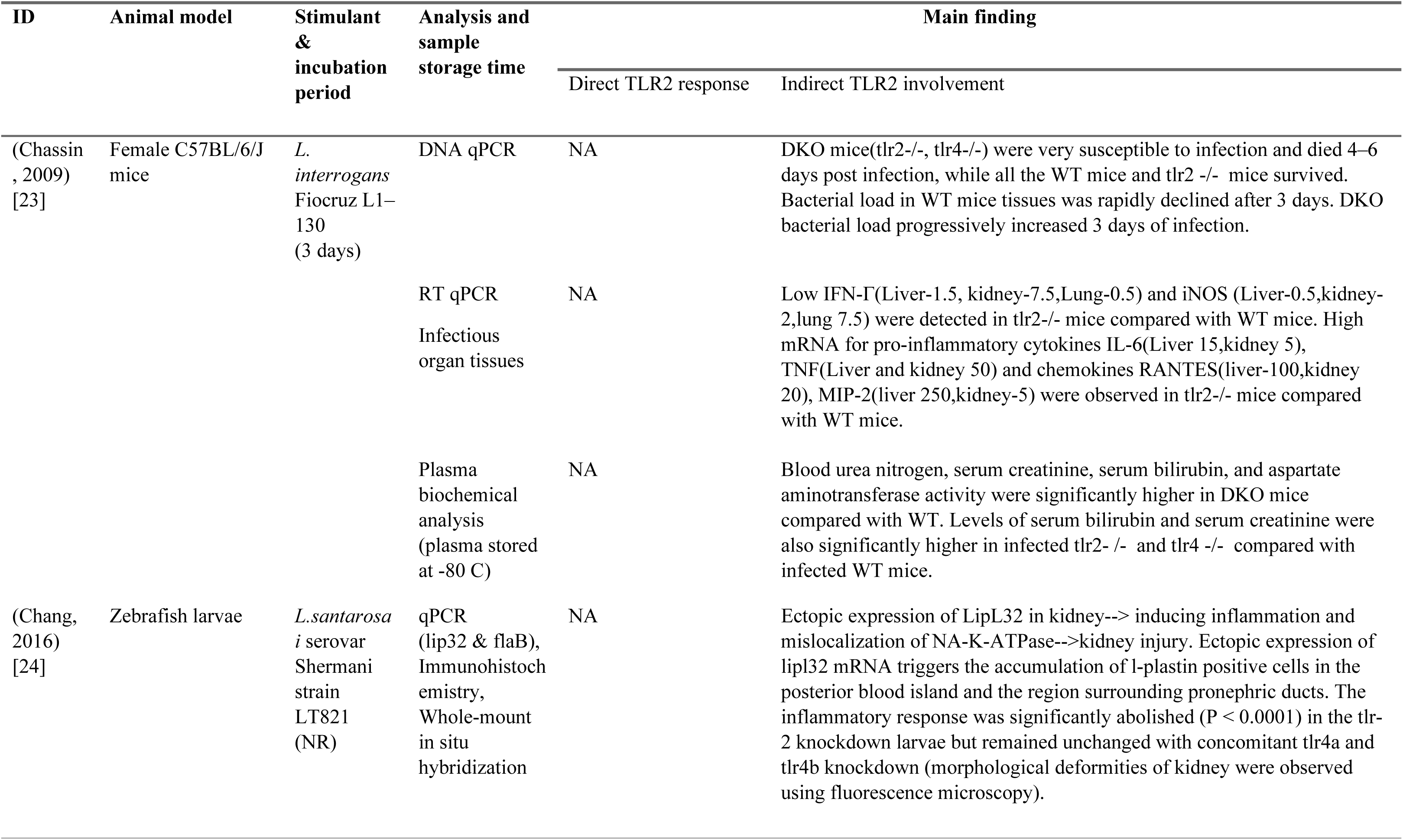
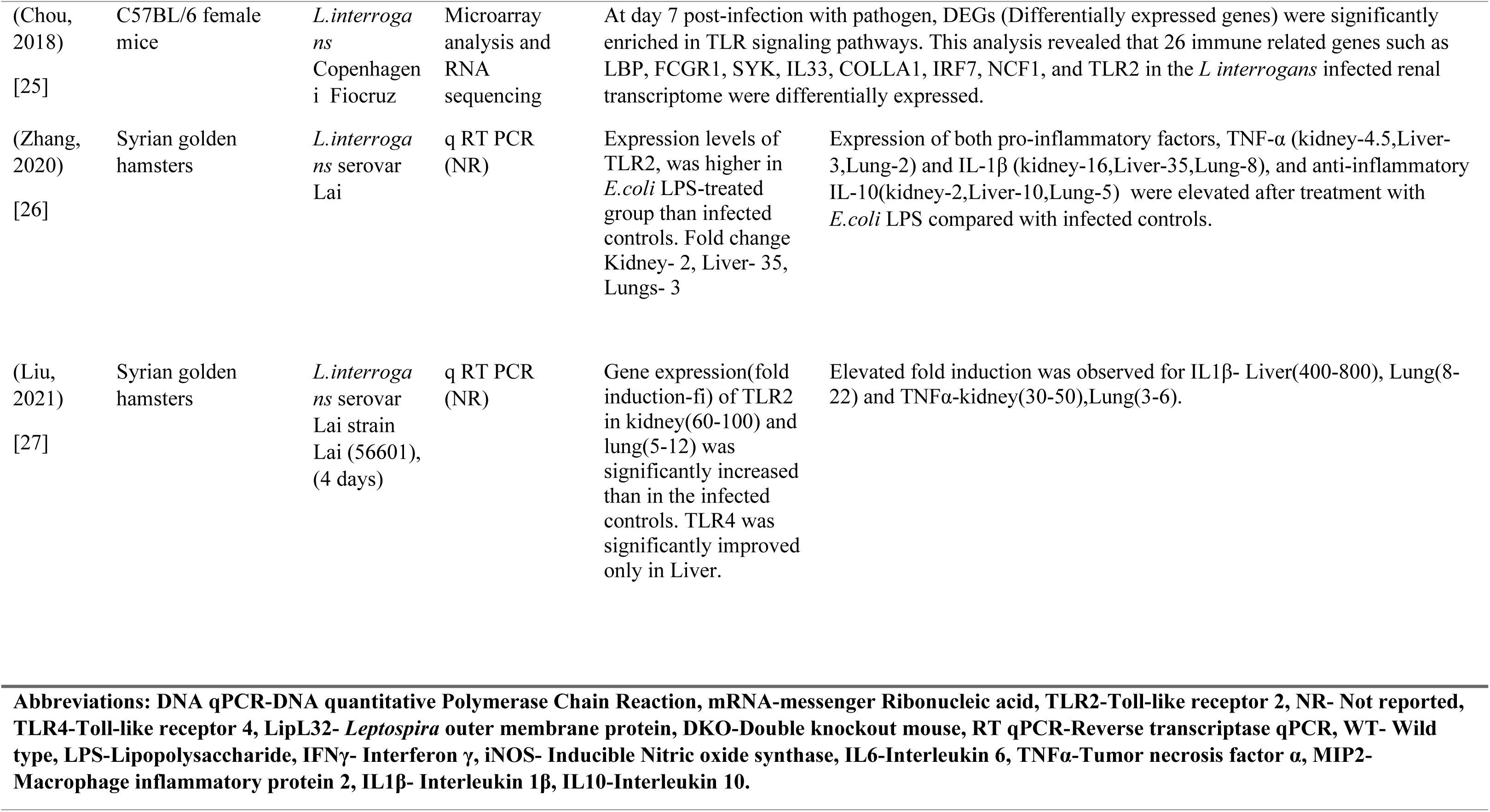
*In-vivo* study characteristics.

### TLR2 response in combined studies of human, *in-vivo* and *in-vitro*

Leptospirosis confirmed human serum samples were used to assess pro and anti-inflammatory cytokines[47] and circulatory micro RNAs[48]. As animal models, mice were used in both studies to assess cytokine levels and micro RNAs during leptospirosis. Infectious agents of mice were *L. interrogans* serogroup Icterohaemorrhagiae Lai[47] and LPS of *L.interrogans* Automnalis strain N2 [48]. THP-1 cells were used in both studies of *in-vitro* experiments. Recombinant hemolysin proteins[47] and LPS of *Leptospira* spp.[48] were used as infectious agents. TLR2 involvement during the pathogenesis response against leptospirosis was observed through the decrease of IL1β,TNFα [47,48], IL6[47], NFκB, IL10[48] secretion/expression due to deficient/knocked down of the TLR2 receptors[47,48]. In addition to TLR2 involvement, TLR4 involvement was also observed during the infection [47]. Table 4 shows the characteristics of combined studies.

**Table 4.**
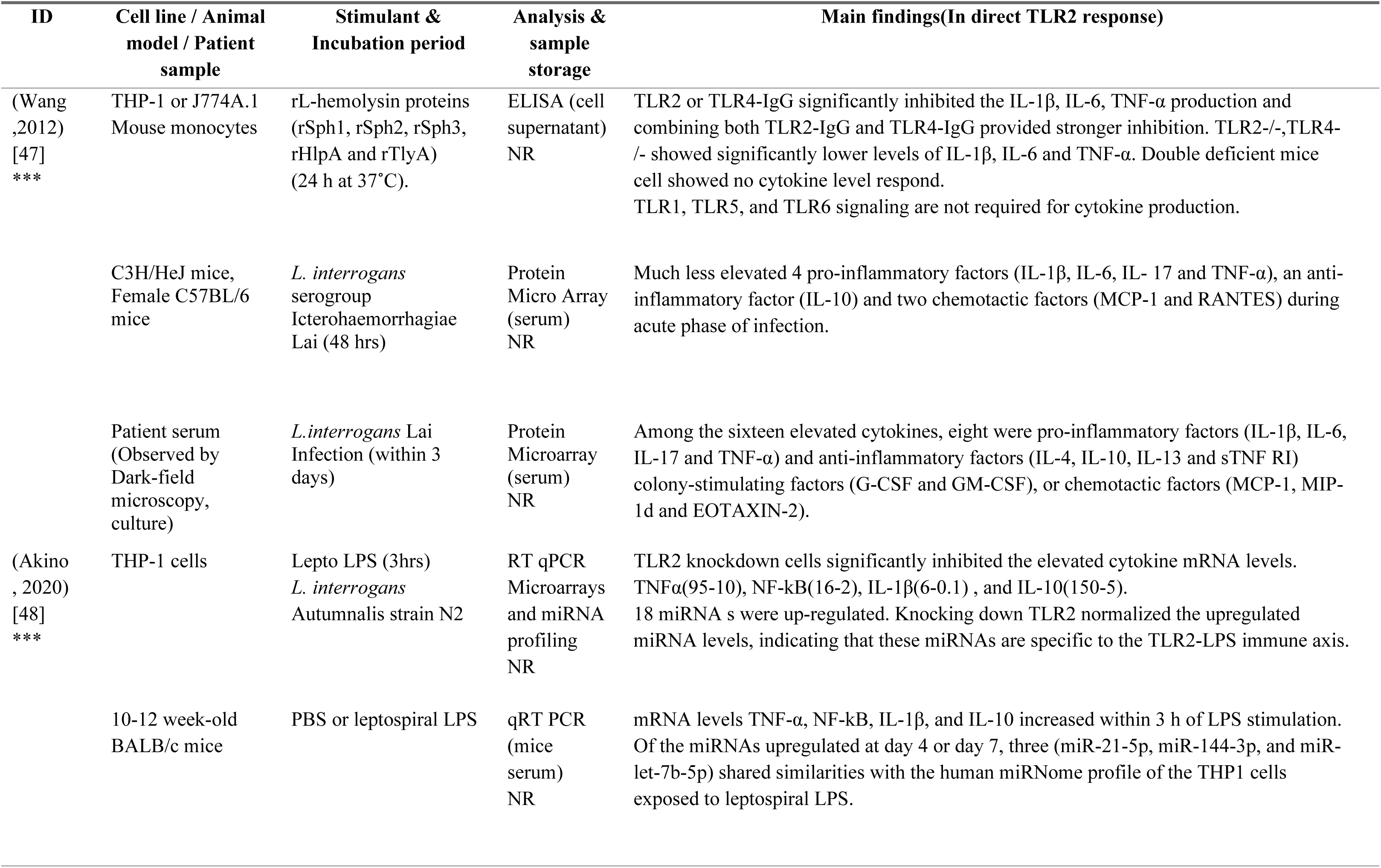
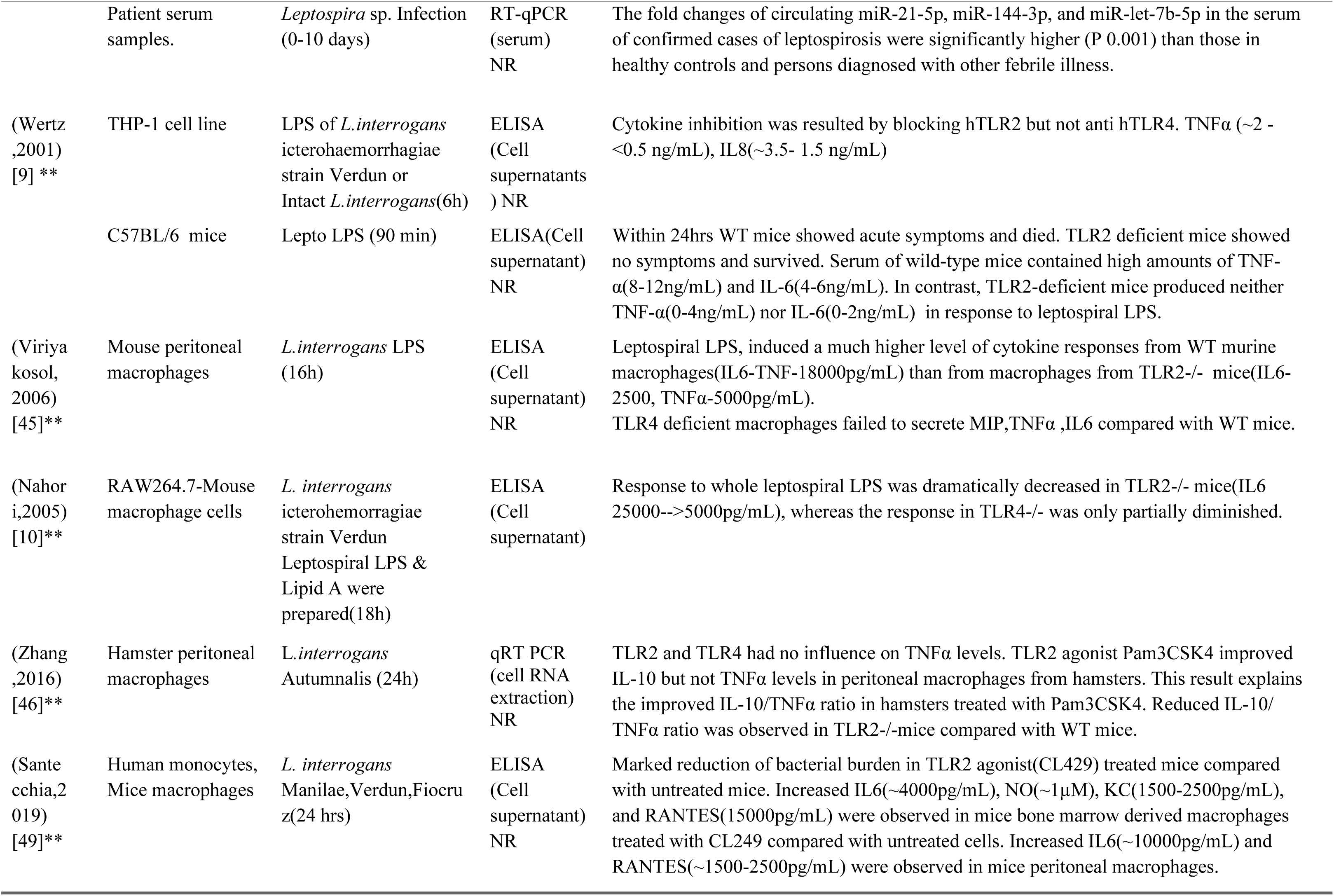
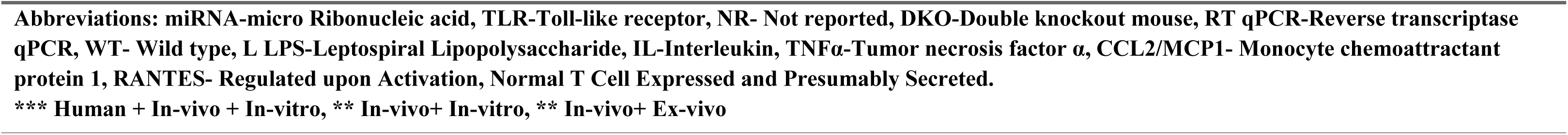
Combined study characteristics.

| ID | Cell line / Animal model / Patient sample | Stimulant & Incubation period | Analysis & sample storage | Main findings(In direct TLR2 response) |
| --- | --- | --- | --- | --- |
| (Wang ,2012) [47] *** | THP-1 or J774A.1 Mouse monocytes | rL-hemolysin proteins (rSph1, rSph2, rSph3, rHlpA and rTlyA) (24 h at 37°C). | ELISA (cell supernatant) NR | TLR2 or TLR4-IgG significantly inhibited the IL-1β, IL-6, TNF-α production and combining both TLR2-IgG and TLR4-IgG provided stronger inhibition. TLR2-/-,TLR4-/- showed significantly lower levels of IL-1β, IL-6 and TNF-α. Double deficient mice cell showed no cytokine level respond. TLR1, TLR5, and TLR6 signaling are not required for cytokine production. |
| | C3H/HeJ mice, Female C57BL/6 mice | *L. interrogans* serogroup Icterohaemorrhagiae Lai (48 hrs) | Protein Micro Array (serum) NR | Much less elevated 4 pro-inflammatory factors (IL-1β, IL-6, IL- 17 and TNF-α), an anti-inflammatory factor (IL-10) and two chemotactic factors (MCP-1 and RANTES) during acute phase of infection. |
| | Patient serum (Observed by Dark-field microscopy, culture) | *L.interrogans* Lai Infection (within 3 days) | Protein Microarray (serum) NR | Among the sixteen elevated cytokines, eight were pro-inflammatory factors (IL-1β, IL-6, IL-17 and TNF-α) and anti-inflammatory factors (IL-4, IL-10, IL-13 and sTNF RI) colony-stimulating factors (G-CSF and GM-CSF), or chemotactic factors (MCP-1, MIP-1d and EOTAXIN-2). |
| (Akino , 2020) [48] *** | THP-1 cells | Lepto LPS (3hrs) *L. interrogans* Autumnalis strain N2 | RT qPCR Microarrays and miRNA profiling NR | TLR2 knockdown cells significantly inhibited the elevated cytokine mRNA levels. TNFα(95-10), NF-kB(16-2), IL-1β(6-0.1) , and IL-10(150-5). 18 miRNA s were up-regulated. Knocking down TLR2 normalized the upregulated miRNA levels, indicating that these miRNAs are specific to the TLR2-LPS immune axis. |
| | 10-12 week-old BALB/c mice | PBS or leptospiral LPS | qRT PCR (mice serum) NR | mRNA levels TNF-α, NF-kB, IL-1β, and IL-10 increased within 3 h of LPS stimulation. Of the miRNAs upregulated at day 4 or day 7, three (miR-21-5p, miR-144-3p, and miR-let-7b-5p) shared similarities with the human miRNome profile of the THP1 cells exposed to leptospiral LPS. |
|  | Patient serum samples. | *Leptospira* sp. Infection (0-10 days) | RT-qPCR (serum)<br>NR | The fold changes of circulating miR-21-5p, miR-144-3p, and miR-let-7b-5p in the serum of confirmed cases of leptospirosis were significantly higher (P 0.001) than those in healthy controls and persons diagnosed with other febrile illness. |
| (Wertz,2001)<br>[9]** | THP-1 cell line | LPS of *L.interrogans* icterohaemorrhagiae strain Verdun or Intact *L.interrogans*(6h) | ELISA (Cell supernatants)<br>) NR | Cytokine inhibition was resulted by blocking hTLR2 but not anti hTLR4. TNFα (~2 - <0.5 ng/mL), IL8(~3.5- 1.5 ng/mL) |
| | C57BL/6 mice | Lepto LPS (90 min) | ELISA(Cell supernatant)<br>NR | Within 24hrs WT mice showed acute symptoms and died. TLR2 deficient mice showed no symptoms and survived. Serum of wild-type mice contained high amounts of TNF-α(8-12ng/mL) and IL-6(4-6ng/mL). In contrast, TLR2-deficient mice produced neither TNF-α(0-4ng/mL) nor IL-6(0-2ng/mL) in response to leptospiral LPS. |
| (Viriya kosol, 2006)<br>[45]** | Mouse peritoneal macrophages | *L.interrogans* LPS (16h) | ELISA (Cell supernatant)<br>NR | Leptospiral LPS, induced a much higher level of cytokine responses from WT murine macrophages(IL6-TNF-18000pg/mL) than from macrophages from TLR2-/- mice(IL6-2500, TNFα-5000pg/mL).<br>TLR4 deficient macrophages failed to secrete MIP,TNFα ,IL6 compared with WT mice. |
| (Nahori,2005)<br>[10]** | RAW264.7-Mouse macrophage cells | *L. interrogans* icterohemorrhagiae strain Verdun<br>Leptospiral LPS & Lipid A were prepared(18h) | ELISA (Cell supernatant) | Response to whole leptospiral LPS was dramatically decreased in TLR2-/- mice(IL6 25000-->5000pg/mL), whereas the response in TLR4-/- was only partially diminished. |
| (Zhang,2016)<br>[46]** | Hamster peritoneal macrophages | *L.interrogans* Autumnalis (24h) | qRT PCR (cell RNA extraction)<br>NR | TLR2 and TLR4 had no influence on TNFα levels. TLR2 agonist Pam3CSK4 improved IL-10 but not TNFα levels in peritoneal macrophages from hamsters. This result explains the improved IL-10/TNFα ratio in hamsters treated with Pam3CSK4. Reduced IL-10/TNFα ratio was observed in TLR2-/-mice compared with WT mice. |
| (Santechia,2019)<br>[49]** | Human monocytes, Mice macrophages | *L. interrogans* Manilae,Verdun,Fiocruz(24 hrs) | ELISA (Cell supernatant)<br>NR | Marked reduction of bacterial burden in TLR2 agonist(CL429) treated mice compared with untreated mice. Increased IL6(~4000pg/mL), NO(~1μM), KC(1500-2500pg/mL), and RANTES(15000pg/mL) were observed in mice bone marrow derived macrophages treated with CL249 compared with untreated cells. Increased IL6(~10000pg/mL) and RANTES(~1500-2500pg/mL) were observed in mice peritoneal macrophages. |

**Abbreviations:** miRNA-micro Ribonucleic acid, TLR-Toll-like receptor, NR- Not reported, DKO-Double knockout mouse, RT qPCR-Reverse transcriptase qPCR, WT- Wild type, L LPS-Leptospiral Lipopolysaccharide, IL-Interleukin, TNFα -Tumor necrosis factor $\alpha$ , CCL2/MCP1- Monocyte chemoattractant protein 1, RANTES- Regulated upon Activation, Normal T Cell Expressed and Presumably Secreted.
**\*\*\* Human + In-vivo + In-vitro, \*\* In-vivo+ In-vitro, \*\* In-vivo+ Ex-vivo**

### TLR2 response in combined studies of *in-vivo* and *in-vitro*

Four studies were combined with *in-vivo* and *in-vitro* experimental models (n=4) (Table 4). The prominent animal model was mice[9,10,45,46] and Syrian golden hamsters[46]. Predominantly used cell lines were mouse macrophages[10,45,46], human peripheral blood monocytes[10], HEK 293[9,10], THP-1[9,10] and CHO-K1 [9]. The mostly used infectious agent was *Leptospira* LPS[9,10]. Other inoculated organisms were *L.interrogans* Icterohaemorrhagiae strain HAI188[45] and *L.interrogans* Autumnalis[46]. TLR2 involvement was observed through inhibition/reduction level of secretion/expression of TNFα[9,45], IL6[9,10,45], IL8[9] and IL10[46] in TLR2 deficient/blocked cells compared with intact cells. TLR4 involvement during the infection was also observed through mediating IL6 and TNFα [10,45]. With regard to survival, LPS treated wild type (WT) mice showed acute illness and death within 24 hours, in contrast TLR2 deficient mice showed no shock and survived[9].

### TLR2 response in combined studies of *ex-vivo* and *in-vivo*

Mice were used as animal models, while mice peritoneal macrophages were used as *ex-vivo* cell line in one study[49] (Table 4). *L.interrogans* Copenhageni and *L.interrogans* Manilae were used as infectious agents. TLR2 Agonist, CL429 treated human monocytes and mice peritoneal cells showed increased level of IL6 and NO, IL1β respectively compared with CL429 untreated cell lines.

### TLR2 response in human studies

Whole blood samples from the confirmed patients with leptospirosis were used in all three human studies (n=3). Microscopic Agglutination Test (MAT) and Polymerase Chain Reaction (PCR) were used as the disease confirmatory test. Significantly increased TLR2 expression observed on polymorphonuclear cells[21] and neutrophils[22] of *Leptospira* infected human whole blood compared with healthy human whole blood. Contrary to that, *Leptospira* infected human monocyte TLR2 expression did not show significant difference compared with healthy human whole blood[20]. In human and *in-vivo* studies, TLR2 response was observed within 7 days of infection, which is longer time period than that observed in *in-vitro* studies (2 days). Table 5 **Error! Reference source not found.**.

**Table 5.** Human study characteristics.

| ID | Leptospirosis confirmation test | Infected organism & incubation period | Analysis & sample storage time period | Main Findings (Direct TLR2 response) |
| --- | --- | --- | --- | --- |
| (Raffray, 2016) [21] | PCR- 23S r RNA<br>IgM- ELISA<br>(>50 U/mL),<br>MAT >1/400 | *L.interrogans*<br>Ictero, Canicola,<br>Ballum<br>(1-7 days) | Flowcytometry<br>(whole blood)<br><6 hrs | Significantly increased (p=0.001) the TLR2 expression on PMN in lepto group (MFI= 0.92) compared with healthy control (MFI=0.57). |
| (Raffray, 2019) [20] | PCR- 23S rRNA<br>IgM-ELISA-<br>(>50 U/mL)<br>MAT >1/400 titer | *L.interrogans*<br>Ictero, Canicola,<br>Ballum<br>(3-5 days) | Flowcytometry<br>(whole blood)<br>NR | The levels of TLR2, TLR4 and CD69 on monocytes were not significantly different between healthy and leptospirosis groups. MFI values; TLR2 (H-10000 L<10000), TLR4 (H~2500 L~2500), CD69 (H~2100 L-2000-2500). |
| (Lindow, 2019) [22] | PCR- 16s rRNA,<br>LipL32<br>MAT > 1/800 | *Leptospira* sp.<br>Infection<br>(1-7 days) | Flowcytometry<br>(whole blood)<br><4hrs | TLR2 expression was significantly higher on neutrophils from patients with more severe (9%) leptospirosis (lung or/& renal dysfunction) & acute(3%)(p<0.05) (lacking organ dysfunction) relative to healthy controls(1%)(p<0.01). (%-percent granulocytes positive for TLR2) |
| **Abbreviations: PCR-Polymerase Chain Reaction, r RNA-ribosomal Ribonucleic acid, IgM-ELISA- Immunoglobulin M-enzyme linked immunosorbent assay, MAT- Microscopic agglutination test, TLR2-Toll-like receptor 2,PMN- Polymorphonuclear, MFI- Mean fluorescence intensity, NR- Not reported, TLR4-Toll-like receptor 4, LipL32- *Leptospira* outer membrane protein.** |  |  |  |  |

### Other TLR response during *Leptospira* infection

Among 32 articles, some of the studies demonstrated the response of TLR4 [10,26–29,45–47] and TLR5 [28] along with the TLR2 response during the infection of *Leptospira* spp./cell components. IL6[10,28,29,47], IL-1β[29,47], TNFα[29,47], MIP[45], CCL2[29], CCL10[29], COX2[29], MCP129), IFNγ[29], iNOS [29] were identified as reduced/down regulated immune mediators in cell lines of deficient/knocked down TLR4 compared with intact cell lines. TNFα reduction was also observed in cells that blocked TLR5(28). Animal studies showed TLR4 involvement in triggering inflammatory mediators such as IL-1β[26,27], TNFα[26,27], IL10[26], NO[26], iNOS[26], anti *Leptospira* IgG[26]. Contribution of both TLR4 and TLR2 in leptospirosis pathogenesis was observed in DKO mice by showing dramatically low mRNA expression of IFNγ and iNOS, compared with WT mice[23].

### Risk of bias in studies

In OHAT risk of bias assessment, all in-vitro and ex-vivo studies (24 of which used cell lines) present a ‘fair’ risk of bias rating. Cell line experiments present a ‘definitely low’ risk for selection bias, but blinding of experimental personnel is ‘not reported’, and performance bias is ‘probably high’. There is ‘definitely low’ risk of bias attrition and detection bias in all studies. All studies present ‘definitely low’ selective reporting bias and other biases. SYRCLE’s risk of bias assessment for 12 in-vivo studies shows a similar rating with an ‘unclear’ risk of bias in 2 domains of selection bias. All studies have an ‘unclear’ risk of performance and detection bias, as there is no report of randomly housed or blinded intervention and no animal random selection for outcome assessment or blinded outcome assessors. No studies have attrition bias, reporting bias, or other biases. In all 5 human studies, NIH-developed quality assessment was used with a similar risk of bias rating. Sample size justification, random selection of patients, and concurrent control selection were not reported, leading to selection and performance bias. Potential confounding variable analyses were not reported, contributing to another risk of bias. Supplementary file S4-S6 Table show the risk of bias results for each study.

### Certainty of evidence

Five domains - risk of bias, inconsistency, indirectness, imprecision, and publication bias - were used to assess the certainty of outcomes in in-vivo and human studies. Risk of bias was assessed and reported for individual studies but not rated down for inherent limitations. Inconsistency was not considered as no meta-analysis was performed, and the direction of estimates did not vary. Indirectness was not down-rated as all studies matched the PICO criteria. Imprecision was down-rated as no studies had sample size justification. Publication bias was not down-rated as no conflicts of interest were present. GRADEpro generated Certainty of the outcome, and ‘increased TLR2 expression during leptospirosis’ in human and in-vivo studies was rated as ‘Moderate’ Table 6 shows certainty of outcomes from *in-vivo* and human studies.

**Table 6.** Certainty of outcomes from *in-vivo* and human studies.

| Study ID | Study design | Risk of bias | Certainty assessment |  |  |  | Impact | Certainty |
| --- | --- | --- | --- | --- | --- | --- | --- | --- |
|  |  |  | Inconsistency | Indirectness | Imprecision | Other considerations |  |  |
| (Chassin,2009)<br>(Chou,2018)<br>[23,25] | observational studies | not serious | not serious | not serious | serious<sup>a</sup> | none | TLR2 involvement against *L.interrogans* infection was observed. | ⊕⊕⊕○<br>Moderate |
| (Zhang,2020)<br>(Liu, 2021)<br>[26,27] | observational studies | not serious | not serious | not serious | serious<sup>a</sup> | none | TLR2 expression was increased compared with infected controls. *E.coli* Lipopolysaccharide and *Iris* polysaccharide treated prior to infection. | ⊕⊕⊕○<br>Moderate |
| (Chang,2016)<br>[24] | observational studies | not serious | not serious | not serious | serious<sup>a</sup> | none | The inflammatory response against *L.santorosai* was significantly abolished in TLR2 knockout zebrafish larvae, but unchanged for TLR4 knockout. | ⊕⊕⊕○<br>Moderate |
| (Raffray 2016)<br>(Lindow,2019)<br>[21,22] | observational studies<sup>a</sup> | not serious | not serious | not serious | serious<sup>b</sup> | none | TLR2 expression was significantly increased on polymorphonuclear neutrophils in the leptospirosis group compared to healthy individuals. | ⊕⊕⊕○<br>Moderate |
| (Raffray,2019)<br>[20] | observational studies<sup>a</sup> | not serious | not serious | not serious | serious<sup>c</sup> | none | No significant difference in TLR2 expression on Monocytes between groups of healthy and leptospirosis. | ⊕⊕⊕○<br>Moderate |

<sup>a</sup>. Not detected specific study design. In terms of experimental design, they used confirmed leptospirosis cases and healthy controls. <sup>b</sup>. Not detected sample size justification. However, a total of 30 patient samples and 17 healthy controls were used. <sup>c</sup>. Not detected sample size justification.

## Discussion

This systematic review presents the response of TLR2 during the infection of pathogenic *Leptospira* spp. or its’ cell components by extracting the data from 32 original studies. As demonstrated by these studies, we have described the response of TLR2 in two approaches; direct TLR2 response and indirect TLR2 involvement through the secretion of cytokine/immune mediators during leptospirosis.

Due to the heterogeneity of the studies, it is challenging to make sense of the role that TLR2 plays during leptospirosis. Nevertheless, some of the narrative review findings are in line with increased TLR2 expression and immune mediator secretion via TLR2. TLR2 activation was observed in mice proximal tubule epithelial cells through *L.santerosai* Shermani inoculation[51,52]. Increased TLR2 expression in mice and hamster models was observed in response to *Leptospira* LipL32 as well[52,53]. Agonist Pam3CSK4 triggered immune response via TLR2, followed by increasing hamster survival rate and reducing organ lesions. Apart from mice/hamster models, pig and bovine cells responded to *Leptospira* LPS via TLR2[54,55]. Aligned with our findings, narrative reviews discussed the renal dysfunction associated with leptospirosis resulted secretion/expression of Fibronectin[56], iNOS[51–53,56,57], CCL2/MCP-1[51,52,56,57], TNFα[51,53,56,57], NFκB[51], CCL2/MIP2[57], CXCL1/KC[57] through TLR2.

A previous systematic review[58] showed elevated level of cytokines (IL-1β, IL-2, IL-4, IL-6, IL-8, IL-10, TNFα) in severe human leptospirosis compared with mild leptospirosis. Since those studies did not explore the TLR2 involvement in triggering immune responses, they did not meet our inclusion criteria. Moreover, similar to our findings, IL-1β, IL-6, IL-8, IL-10, TNFα cytokines showed elevation in severe human leptospirosis. As per our findings, recognition of PAMPs by the host TLR2 triggers the activation of immune effectors such as cytokines/chemokines IL6[9,10,28,29,37,41,45,47,49], IL8[9,38,41–43], IL 1β[29,41,43,47–49], TNFα[9,28,29,37,38,41,43,45,47,48], IFNγ[29], IL10[46], CCL2/MCP-1[29,31,33,38,39], CCL10[29,38–40], COX2[29], CXCL1/KC[39], and CXCL2/MIP2[40] followed by leukocyte recruitment to heal tissues/organ lesions. While clearing bacteria, antimicrobial peptides hBD2[41,43], iNOS[31,33], Fibronectin[30], and Oxygen and Nitrogen reactive species are produced[23,26]. Although inflammatory response is defensive against the bacteria, excessive levels can cause harmful effects such as organ lesions[24,46]. To determine whether innate TLR2 immunity has a protective or harmful effect, it is crucial to recognize the level and threshold of pathogen-host immune responses, thereby accurately interpreting the role of TLR2. As per our results, TLR2 activator (Agonist) and the TLR2 blocker (Antagonist) can be used to modulate the level of immune mediators and enhance the immune response against the infection. Another crucial factor in identifying the TLR2 response is the incubation period of the bacterium in the host tissues or host body. *In-vitro* studies observed TLR2 mRNA expression within 48hrs of infection, while human and *in-vivo* studies showed TLR2 response no longer than 7 days after the infection.

The main limitation observed is the limited number of human studies identified during the search. While answering the question on the role of TLR2 during leptospirosis makes it challenging to anticipate how accurately these findings reflect in humans due to the utilization of diverse cells and experimental models in *in-vitro* and *in-vivo* studies. It is worth to know the exact time period of TLR2 expression after the onset of *Leptospira* exposure that was lacking in most of the studies. Another limitation presents with inherent qualities of *in-vivo* studies; not reporting of allocation sequence generation, allocation concealment, animal randomization, and blinding of caregivers and assessors led to rate down the quality of the studies. Concerning the strengths, even with the inherent study qualities, human, *in-vivo*, *in-vitro*, and *ex-vivo* studies present an overall risk of bias rating as ‘Fair’, ‘Fair’, ‘Good’, and ‘Good’, respectively. GRADE approach assessed the important outcomes of *in-vivo* and human studies as ‘Moderate’ certainty. To the best of our knowledge, this study is the first systematic review that address the TLR2 response during leptospirosis.

## Conclusions

The scarcity of human studies hinders the establishment of a robust evidence base for interpreting the direct response of human Toll-like receptor 2 (TLR2) during leptospirosis. However, it has been observed that increased TLR2 expression and the secretion/mRNA expression of various cytokines/chemokines (IL6, IL8, IL 1β, TNFα, IFNγ, IL10, CCL2/MCP-1, CCL10, COX2, CXCL1/KC, CXCL2/MIP2) and immune effectors (antimicrobial peptides hBD2, iNOS, Fibronectin, Oxygen, and Nitrogen reactive species) are significant functions of host TLR2 in leptospirosis. While an immune response against the bacterium is crucial for overcoming the disease, the subsequent detrimental effects of excessive immune mediators on tissues/organs cannot be ignored. The findings of this systematic review provide valuable insights for the development of new therapeutic strategies aimed at maintaining a moderate level of immune mediators through the use of TLR2 receptor agonists or antagonists. Furthermore, indirect involvement of host TLR4 and TLR5 through the secretion of immune mediators was also identified in leptospirosis.

## Other Information

### Registration and protocol

International prospective register of systematic reviews. PROSPERO 2022 CRD42022307480 (No amendments were provided at registration)

Available from https://www.crd.york.ac.uk/prospero/display_record.php?ID=CRD42022307480

Protocol was not prepared.

### Supporting information

**S1. Search strategies for the systematic review**

**S2. PRISMA 2020 Abstract check list**

**S3. PRISMA 2020 check list S4. Risk of bias Assessment**

## Availability of data, code and other materials

All the data produced or examined during this study have been incorporated in the published article (along with supplementary materials).

